# N-cadherin-Presented Slit Repulsive-Cues Direct Collective Schwann cell Migration

**DOI:** 10.1101/549030

**Authors:** Julian J.A Hoving, Elizabeth Harford-Wright, Patrick Wingfield-Digby, Anne-Laure Cattin, Mariana Campana, Toby Morgan, Victor Quereda, Erica Torchiaro, Alison C. Lloyd

## Abstract

Collective cell migration is fundamental for the development of organisms and in the adult, for tissue regeneration and in pathological conditions such as cancer. Migration as a coherent group requires the maintenance of cell-cell interactions, while contact-inhibition-of-locomotion (CIL), a local repulsive force, propels the group forward. Here we show that the cell-cell interaction molecule, N-cadherin, regulates both adhesion and repulsion processes during Schwann cell collective migration, which is required for peripheral nerve regeneration. However, distinct from its role in cell-cell adhesion, the repulsion process is independent of N-cadherin trans-homodimerisation and the associated adherens junction complex. Rather, the extracellular domain of N-cadherin acts to traffic a repulsive Slit2/Slit3 signal to the cell-surface. Inhibiting Slit2/Slit3 signalling inhibits CIL and subsequently collective SC migration, resulting in adherent, non-migratory cell clusters. These findings provide insight into how opposing signals can mediate collective cell migration and how CIL pathways are promising targets for inhibiting pathological cell migration.

## Introduction

Tissue and organ morphogenesis requires the orchestration of the movement of large numbers of cells^1, 2^. In the adult, cell migration is less frequent but is important for aspects of tissue renewal and immune surveillance and can be activated following an injury, to contribute to wound healing and tissue-regeneration^3–6^. Moreover, these modes of migration are frequently recapitulated during pathologies such as cancer, allowing tumour cells to spread from their original site^3, 7, 8^.

Peripheral nerve is one of the few tissues in the adult that retains the remarkable ability to regenerate following an injury^9–11^. We have previously shown that the successful regeneration of a transected nerve requires the collective migration of cords of Schwann cells (SCs) to transport regrowing axons the injury site ^12^. Moreover, SC cords gain directionality across the wound, by migrating along a newly formed polarised vasculature, which develops prior to SC migration^13, 14^. However, SCs cultured alone exhibit contact inhibition of locomotion (CIL)^12^, a process where two cells repulse each other upon contact as a result of the inhibition of local protrusion formation and the repolarisation of the cells towards an alternative migration path^15–18^. Following an injury however, SC collective-migration is triggered following interactions with fibroblasts that come into contact with SCs as they enter the wound-site^12^. This heterotypic interaction with fibroblasts transforms SC behaviour from repulsive to attractive, a process mediated by ephrinB/EphB2-signalling inducing a Sox2-mediated re-localisation of N-cadherin to the site of cell-cell junctions, resulting in their migration as cellular cords^12^. Consistent with this, we showed that in mice lacking EphB2 signalling (*EphB2^-/-^*) both SC migration and axonal regeneration are impaired.

Recently, CIL has been shown to play a role in the dispersal of cells during development, with CIL promoting the spread of cells through tissues^19, 20^. Moreover, CIL is also important for providing an outward force during collective cell migration, in that, similarly as seen with single cells, CIL regulates the polarisation of cells, by inhibiting local protrusions at the site of cell-cell contact and inducing protrusion formation at the free edge, thus promoting outward migration^15–18, 21, 22^. This implies that collective migration requires the maintenance of a CIL signal in the presence of a stronger adhesive signal, but how this can be achieved is poorly understood.

Here we show that N-cadherin mediates both the adhesion and repulsive forces required for collective SC migration but via two distinct mechanisms; while adhesion is dependent on the Sox2-stabilised, N-cadherin adherens-junction complex, repulsion is the result of N-cadherin presenting a Slit2 and Slit3 repulsive signal. Moreover, inhibiting the CIL signal results in tight clusters of non-migratory cells. The ability of N-cadherin to simultaneously regulate adhesion and CIL shows how these opposing processes can be coordinated during collective migration and that CIL signals are an attractive target for inhibiting unwanted collective cell migration.

## Results

### N-cadherin is required for contact inhibition of locomotion between Schwann cells

We previously showed that EphB2 activation of Sox2 results in the clustering of SCs but video analysis showed these clustered cells appeared to maintain CIL, as the cells appeared to be attempting to migrate away from the cluster^12^. This is consistent with the migration of SC cords during nerve regeneration, which would be predicted to require a force such as CIL to drive migration forward^18, 21, 23, 24^. N-cadherin has been implicated in CIL in other cell types^21, 22, 25^, so we addressed whether N-cadherin was also required for the regulation of collective SC migration. To do this, we initially blocked N-cadherin expression using two independent siRNAs (Fig. 1a-b), and performed a wound-healing assay (Fig. 1c-e). Time-lapse microscopy showed that while scrambled-treated cells migrated in a directional manner to efficiently close the gap, N-cadherin-knockdown cells closed the gap more slowly, as they migrated in multiple directions, over each other and with a lack of persistent migration towards the gap (Fig. 1c-d and Supplementary Fig. 1a). This difference was not due to a defect in migration speed, as individual N-cadherin-knockdown cells migrated more rapidly (Fig. 1e), suggesting that N-cadherin was required for the cell-contact dependent process driving outward migration. Consistent with this, confluent control SCs form a monolayer and stable junctions, whereas N-cadherin knockdown cells were unable to form junctions and grew on top of each other (Fig. 1b and Supplementary Fig. 1b), suggesting they have lost the ability to recognise each other, and that the loss of contact-dependent outward migration observed in the wound-healing assay, could be due to loss of CIL.

**Figure 1.**
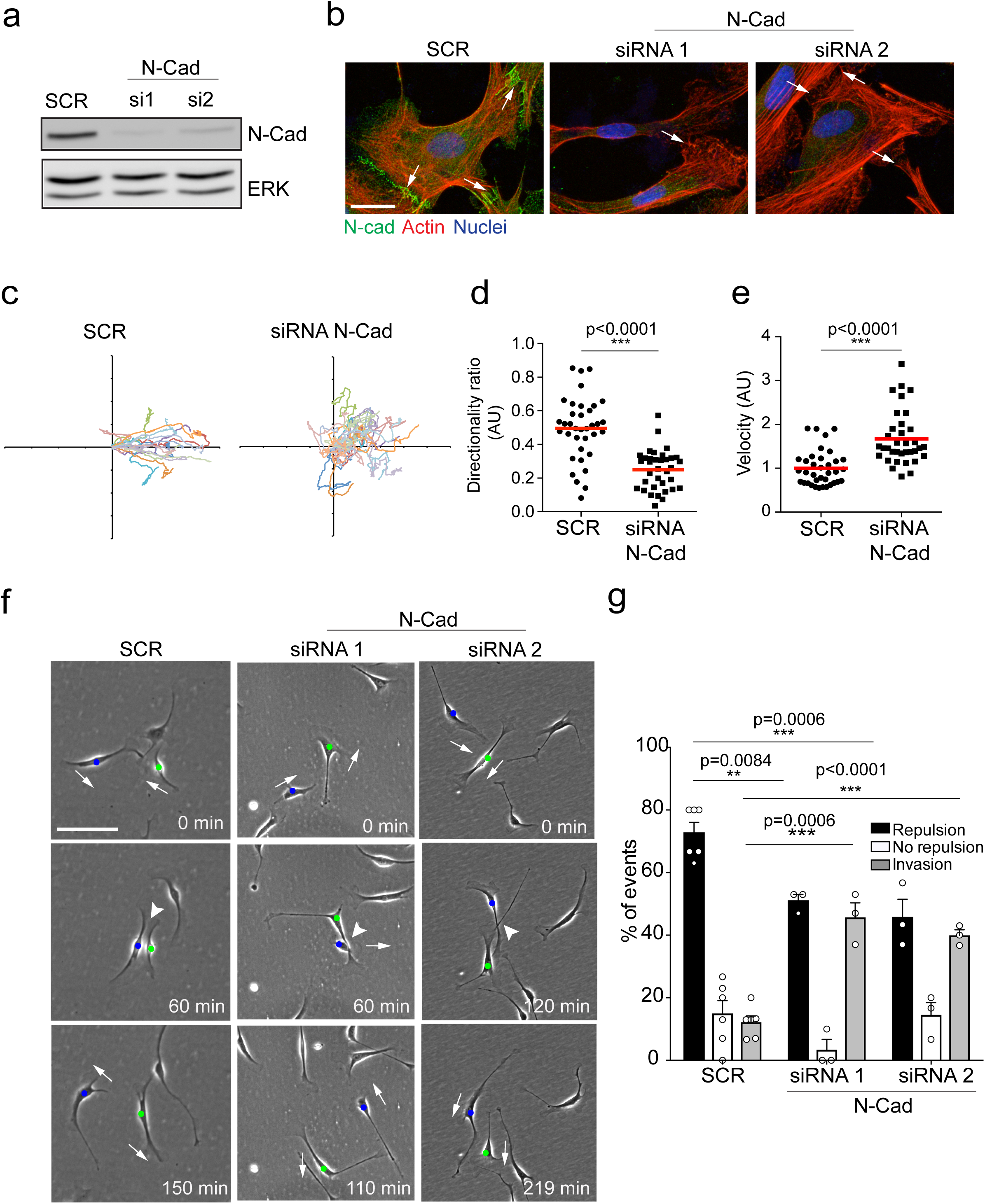
N-cadherin is required for contact inhibition of locomotion between Schwann cells. **a.** Representative western blot of three independent experiments showing N-cadherin (N-cad) protein levels in SCs treated with either 2nM Scrambled (SCR), siRNA1 (si1) or siRNA2 (si2) for 48 hours. ERK was used as a loading control. **b.** Representative confocal images of scrambled, siRNA1 or siRNA2 treated cells at 36 hours, immunostained to detect N-cad (green), Actin (red) and the nuclei (blue). White arrows indicate the cell contacts. Scale bar=20µM. **c.** A representative graph of three independent experiments showing the trajectories of scrambled treated cells or N-cad knockdown cells treated with siRNA1. n= 36 and 35 for scrambled and N-cad knockdown cells respectively. **d-e.** Quantification of the directionality ratio **(d)** and the velocity **(e)** from the cells tracked in **(c)**. The red line indicates the mean. ***p<0.001 compared to SCR. **f.** Representative time-lapse images of a CIL assay, showing scrambled or N-cad knockdown cells, treated with siRNA1 or siRNA2 that repulsed or invaded respectively (arrow heads). The blue and green dots indicate the two interacting cells. Arrows indicate the direction of migration. Scale bar=100µm. **g.** Quantification of (**f)** (mean ± SEM) n=3. Note, that random protrusions (SCR (n=36) NCAD (35 cells) and retractions are produced by SCs in all directions that leads to a high number of apparent repulsion events, even when loss of CIL has occurred. **p<0.01, ***p<0.001.

To study the role of N-cadherin in homotypic CIL between SCs, we quantified CIL upon cell-cell contact of SCs cultured at low density. Live imaging of scrambled-treated SCs, showed that SCs migrate in multiple directions, making frequent, seemingly random, protrusions and retractions in multiple directions. However, upon contact with another SC, a SC retracts the contacting protrusion, appears repulsed, and changes its direction of migration (Fig. 1f, quantified in Fig. 1g), behaviours that are features of CIL^15, 16^ and is consistent with what we observed previously^12^. In contrast, N-cadherin knockdown cells have lost the repulsion signal, meaning that protrusions of N-cadherin knockdown cells continued to move forward upon contact, resulting in the migration of cells on top of each other, a behaviour we termed invasion (Fig. 1f, quantified in Fig. 1g). These results show that in addition to mediating adhesion between SCs, N-cadherin is also required for CIL between collectively migrating SCs.

### CIL is independent of the adherens junction complex

To understand how N-cadherin can mediate both cell-cell adhesion and CIL, we addressed whether both processes act via the N-cadherin adhesion complex, which has previously been implicated in both processes^12, 21, 22^. N-cadherin transmits adhesive forces between neighbouring cells by forming trans-homodimers which relay signalling via a well-characterised intracellular adhesion complex to the actin cytoskeleton^26, 27^. To test this, we initially used time-lapse microscopy to analyse CIL between red-labelled, scrambled-treated cells and green-labelled, N-cadherin knockdown cells in co-cultures to determine the requirement for N-cadherin homodimers between the cells (Fig. 2a). Surprisingly, while an N-cadherin knockdown cell (e.g. cell N2) invaded another N-cadherin knockdown cell (N1), the same cell was repulsed upon subsequent contact with a scrambled-treated cell (S2) (Fig. 2a, quantified in Fig. 2b). This suggested that N-cadherin is required to present a repulsion signal and induce repulsion, but is not required for a cell to be repulsed. Consistent with this, when analysing the response of the scrambled-treated cells (e.g. S2) upon contact with an N-cadherin knockdown cell (e.g. N2), the majority of the scrambled-treated cells were not repulsed (Fig. 2a, quantified in 2b). However, the scrambled-treated cells did not invade the N-cadherin knockdown cells, as the N-cadherin knockdown cells were repulsed and migrated away. This showed that N-cadherin is required to present a repulsion signal and induce repulsion, but not to be repulsed.

**Figure 2.**
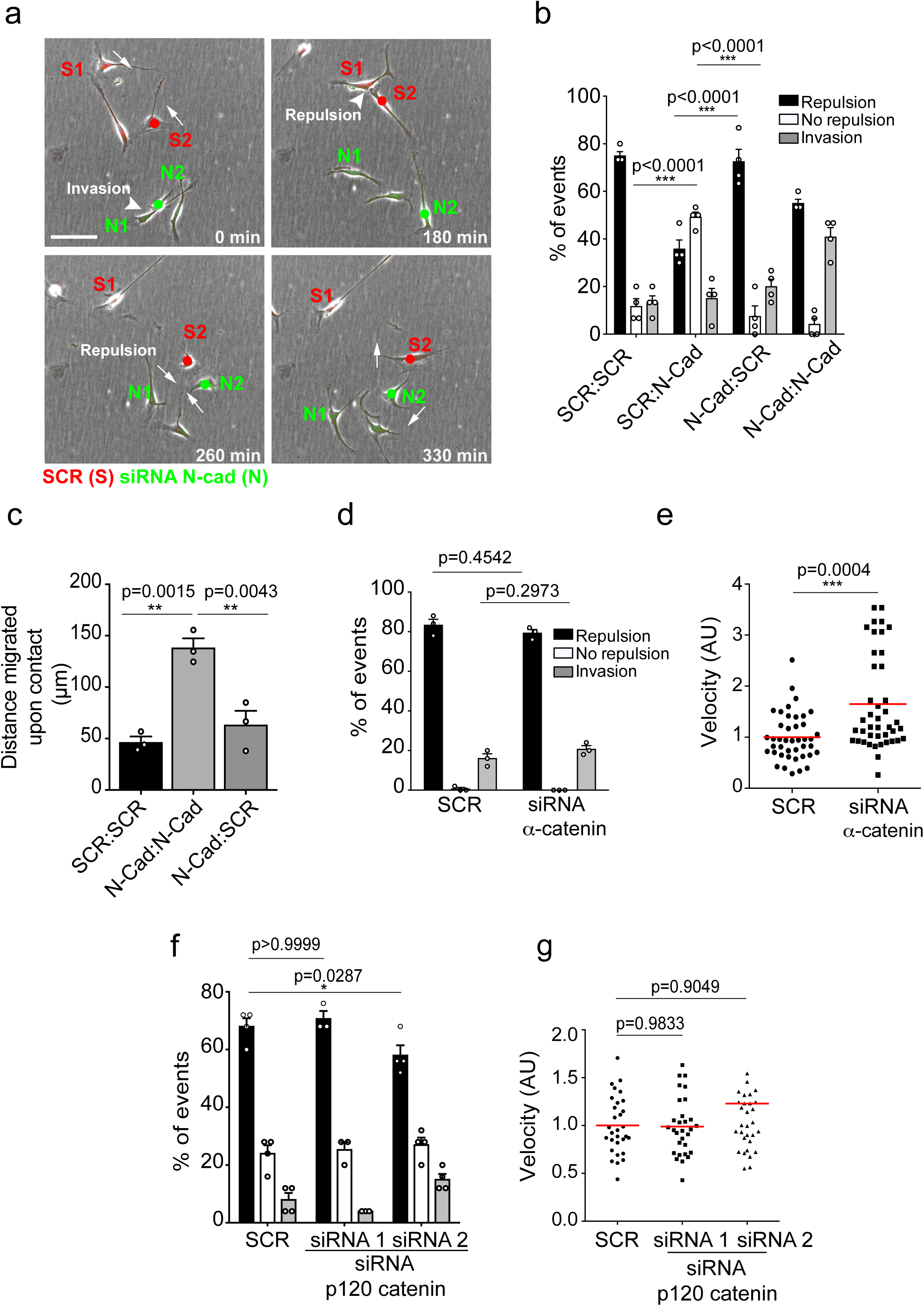
CIL is independent of the adherens junction complex. **a.** Representative time-lapse microscopy images of a CIL assay in which red fluorescence-labelled scrambled-treated cells (S1 and S2) were mixed with green fluorescent-labelled N-cad knockdown cells (N1 and N2). Cells of interest are indicated with a red or green dot for scrambled and N-cad knockdown cells, respectively. Arrows indicate direction of migration. Invasion and repulsion events are indicated. Scale bar=100µm. **b.** Quantification of **(a)** (mean ± SEM) n=3 ***p<0.001. **c.** Quantification of the distance migrated upon contact in scrambled, N-cad knockdown or scrambled and N-cad knockdown cells seeded in chambers. (mean ± SEM) n=3. **p<0.01. **d-e.** Quantification of CIL and velocity in scrambled and α-catenin knockdown cells (mean ± SEM) n=3. ***p<0.001. **f-g.** Graphs show the quantification of CIL and velocity of the tracks from scrambled treated cells (dots) and p120-catenin knockdown cells treated with siRNA1 (squares) or siRNA2 (triangles). Red lines indicate the mean. n=3. *p<0.05.

To further test this observation, we performed an invasion assay, which assessed the ability of cells to form a boundary upon gap-closure in a wounding assay. Consistent with the single-cell analysis, we found that whereas control cells were able to form a boundary upon wound closure and N-cadherin knockdown cells invaded other, N-cadherin knockdown cells were repulsed by the approaching scrambled-treated cells and so were unable to invade (Fig. 2c and Supplementary Fig. 2a). These results confirmed that N-cadherin is only required to present the repulsive CIL signal. Moreover, it suggests that N-cadherin mediates CIL independent of trans-homodimerisation, implying a distinct mechanism mediates CIL.

While the repulsion signal is independent of the trans-homodimerisation of N-cadherin, it remained a possibility that the classical adhesion molecules associated with N-cadherin were required for the CIL signal. We therefore tested whether α-catenin and p120-catenin, which are known to physically connect N-cadherin to the actin cytoskeleton and regulate the stability of N-cadherin at the cell-surface respectively, are required for CIL^28–33^. Time-lapse microscopy showed that efficient knockdown of α-catenin in SCs (Supplementary Fig. 2b) had no effect on CIL, as the cells still repulsed each other despite an increase in cell velocity (Fig. 2d-e). However, consistent with previous reports, the connection of N-cadherin to the actin cytoskeleton appeared to be disrupted, as shown by the more cortical appearance of the actin cytoskeleton at N-cadherin cell-cell contacts (Supplementary Fig. 2d). Similarly, SCs depleted of p120 catenin (Supplementary Fig. 2c) were still repulsed upon contact, although there was a slight but significant decrease in repulsion in cells treated with siRNA2 (Fig. 2f). Consistent with the role of p120-catenin in regulating cadherin levels at the cell-surface^30, 34–36^, we found a strong decrease in the levels of N-cadherin at the cell surface, although this was still concentrated at cell-cell contacts, and this was greater in cells treated with the more efficient siRNA2, making it likely that a decrease in N-cadherin at the cell-surface was responsible for the small effect on CIL (Fig. 2f-g and Supplementary Fig. 2e). Together these results indicated that N-cadherin-mediated CIL is both independent of trans-homodimerisation and acts independently of the adherens junction complex.

### The extracellular domain of N-cadherin is sufficient to mediate CIL

To investigate how N-cadherin mediates CIL, we tested which domains of N-cadherin were sufficient to rescue CIL in N-cadherin knocked-down cells. Western blotting showed that either full length tomato-tagged N-cadherin or mutants, that either lack the intracellular domain, or the extracellular domain were expressed at similar levels (Fig. 3a-b) ^37^. Moreover, the constructs behaved as predicted; confocal images showed that full-length exogenous N-cadherin was localised at cell-cell junctions, and co-localised with α-catenin and p120-catenin. In contrast, the intracellular domain of N-cadherin localised at the membrane, but did not form junctions, but was able to recruit α-catenin and p120 catenin, whereas, the extracellular domain of N-cadherin formed junctions but was unable to recruit α-catenin and p120-catenin (Supplementary Fig. 3).

**Figure 3.**
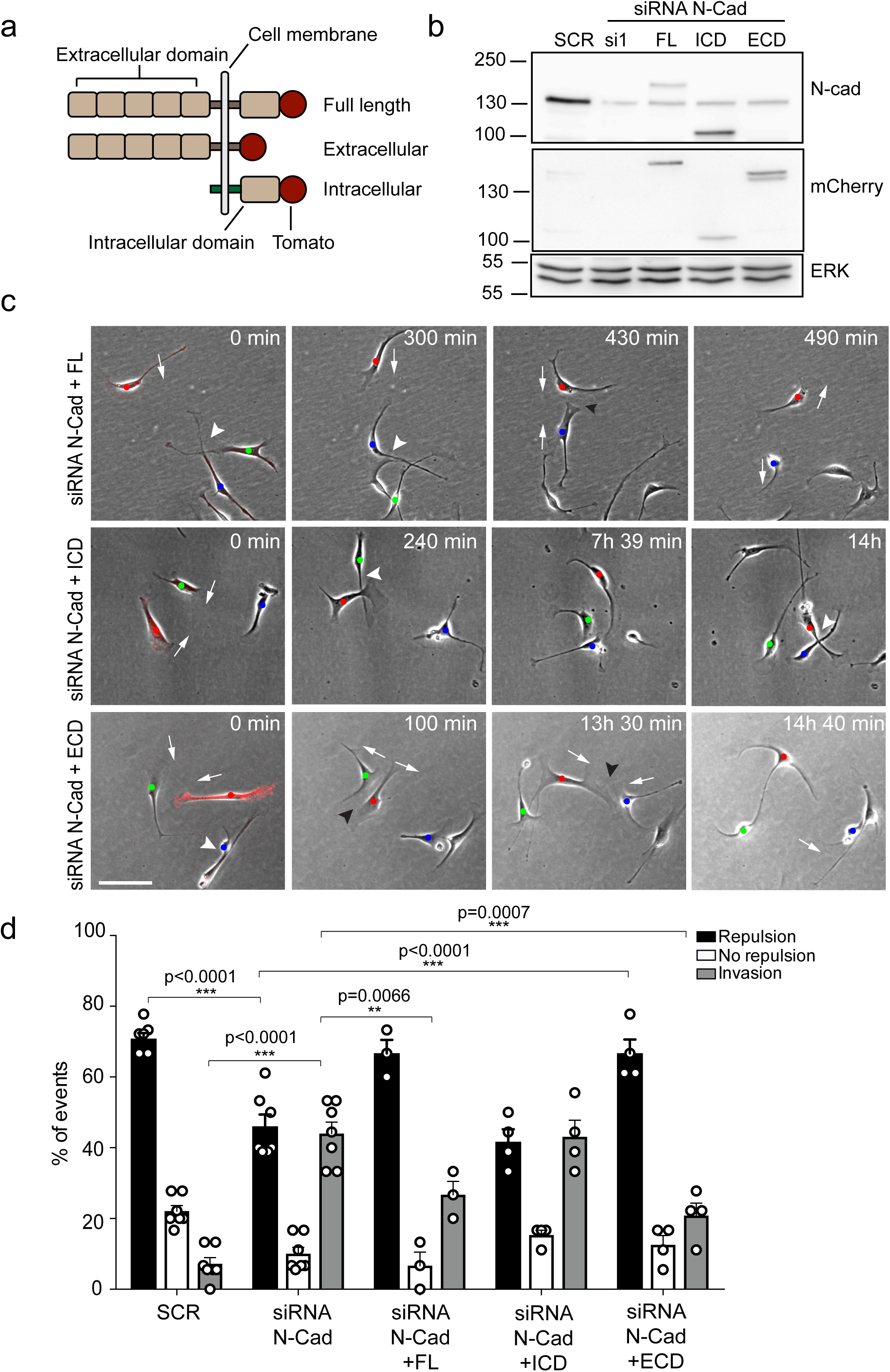
The extracellular domain of N-cadherin is sufficient to mediate CIL. **a**. Schematic of N-cad full length, extracellular and intracellular domains tagged with ptdTomato at the C-terminus. The intracellular domain of N-cad has an additional Lyn membrane-targeting sequence at the N-terminus to target it to the membrane. **b.** Representative Western blot using antibodies that recognise the C-terminus of N-cad and mCherry showing the expression levels of the constructs, 48 hours after knockdown of endogenous N-cad using siRNA1. ERK was used a loading control. **c.** Representative time-lapse microscopy images from a CIL assay of N-cad knockdown cells transfected with the full-length of N-cad, the intracellular domain of N-cad (siRNA1+ICD) or the extracellular domain of N-cad (siRNA1+ECD) tagged with ptdTomato. Arrows indicate the direction of migration. Black arrowheads indicate repulsion events (siRNA1+ full length and ECD). White arrowheads indicate invasion events (siRNA1 + Full length, ICD and ECD). Cells of interest that are interacting are indicated by blue, red, and green dots. Scale bar=100µm. **d.** Quantification of **(c)** Full length (n=3), ECD and ICD (n=4), SCR and siRNA1 (n=7) (mean ± SEM) **p<0.01, ***p<0.001.

To analyse the effect of these constructs on CIL, we quantified the response of N-cadherin knockdown cells upon contact with a N-cadherin construct-expressing cell at low density, as we showed earlier that N-cadherin knockdown cells were still repulsed by an N-cadherin expressing cell. As expected, while N-cadherin knockdown cells invaded each other, they were repulsed upon contact with cells rescued with full-length N-cadherin (Fig. 3c, quantified in Fig. 3d), confirming that loss of CIL is specific to N-cadherin. In contrast, CIL was not rescued by SCs expressing the intracellular domain of N-cadherin (Fig. 3c-d). Strikingly, however, CIL was fully rescued by SCs expressing the extracellular domain of N-cadherin showing that intracellular signalling by N-cadherin is not required to mediate CIL (Fig. 3c-d). This result is consistent with our findings that the adherens junction complex is not required to mediate N-cadherin mediated CIL and together with our findings that N-cadherin is only required to present a repulsion signal suggests an additional co-repulsion signal may be required.

### Glypican-4 and Slit2/3 are required for CIL

We have previously shown that SCs are repulsed by fibroblasts in an ephrinB/EphB2-dependent manner^12^, followed by cell clustering due to the activation of Sox2-dependent relocalisation of N-cadherin to cell-cell junctions. We thus reasoned that ephrinB/EphB2 was unlikely to be responsible for the homotypic CIL between SCs and this was confirmed in knockdown experiments (Supplementary Fig. 4a).

In an attempt to identify the homotypic CIL signal, we performed a series of proteomic screens, using N-cadherin or the extracellular domain of N-cadherin as bait, followed by co-immunoprecipitation and mass spectrometry analysis (data not shown). In one of these analyses, peptides of Glypican-4, a Glycosylphosphatidylinositol (GPI)-linked heparan sulfate proteoglycan, were specifically detected in N-cadherin pull-down samples. Glypicans have previously been shown to play a role in axonal guidance and collective cell migration, so although not implicated in CIL were an interesting candidate^38–40^. To test if Glypican-4 was required for mediating repulsive signals between SCs, we knocked down Glypican-4 and performed a CIL assay (Supplementary Fig. 4b). Intriguingly, Glypican-4 knockdown cells did not repulse upon contact nor did they invade (Fig. 4a, quantified in 4b), rather, it appeared that the cells upon contact adhered to each other, causing the cells to form clusters (Fig. 4c, quantified in 4d) without affecting cell velocity prior to cluster formation (Supplementary Fig. 4c). This result was in stark contrast to N-cadherin knockdown cells, which invaded upon contact (Fig. 1 f-g). Moreover, the cell-clusters behaved differently to Sox2 expressing cells in that upon contact, they no longer appeared to show repulsion, resulting in “quieter” clusters that were no longer polarised towards outward migration (Figure 4c). Consistent with this finding, N-cadherin was still present at the membrane in Glypican-4 knockdown cells with confocal images showing that N-cadherin accumulated at the cell-cell contacts, forming longer junctions compared to the control (Fig. 4c). This suggested that Glypican-4 is involved in the SC CIL signal and that its absence (because N-cadherin remains at the cell surface) results in the formation of more stable homotypic adhesions, resulting in non-polarised clusters of cells.

**Figure 4.**
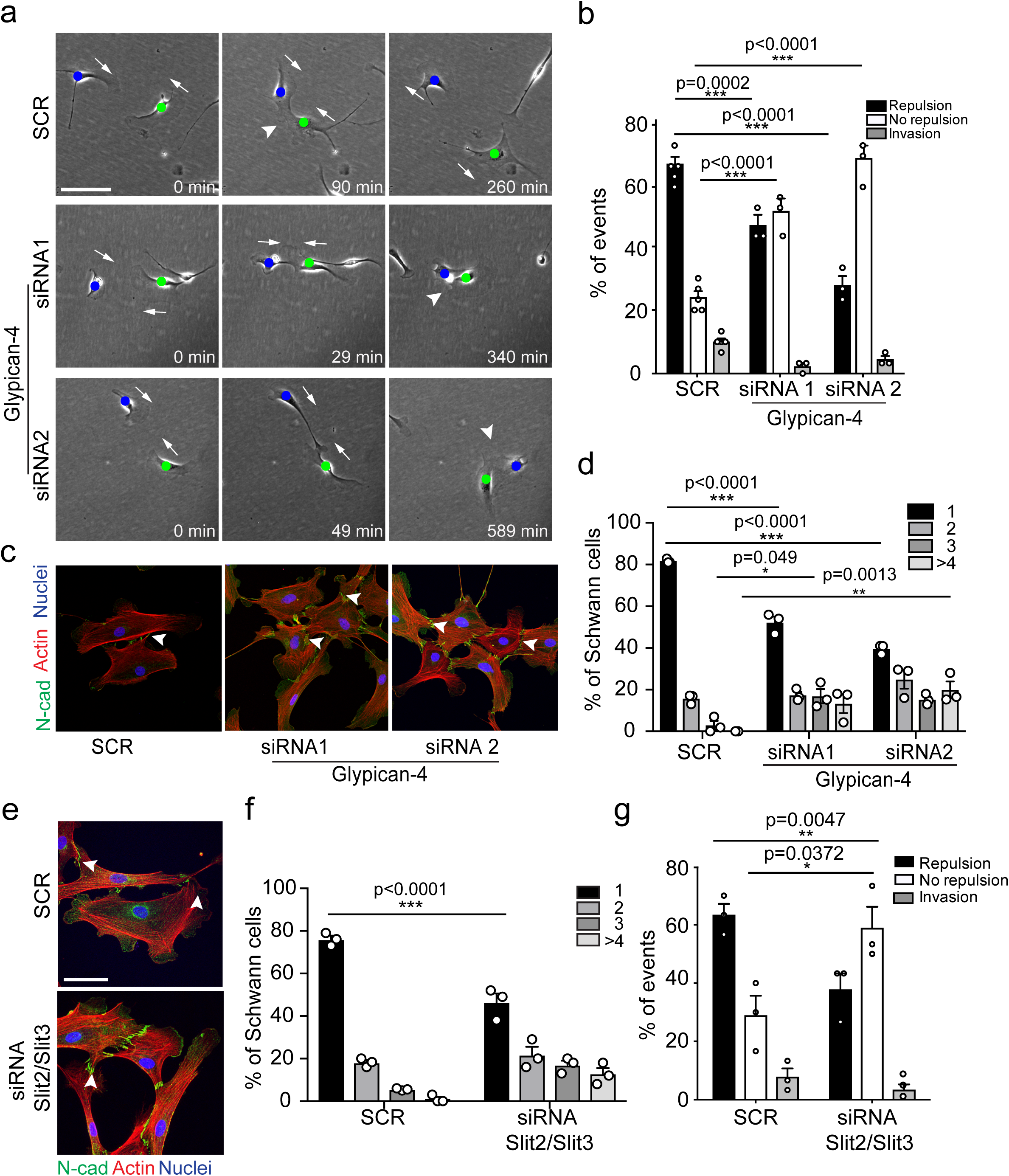
Glypican-4 and Slit2/3 are required for CIL. **a.** Representative time-lapse images of a CIL assay, showing scrambled or glypican-4 knockdown cells, treated with siRNA1 or siRNA2, that are repulsed or not repulsed upon contact respectively (arrow heads). Arrows indicate direction of migration. The green and blue dots indicate the interacting cells. Scale bar=100µm **b.** Quantification of **(a)** (mean ± SEM) n=3. ***p<0.001. **c.** Representative confocal images showing scrambled or glypican-4 knockdown cells stained with Actin (red) and N-cad (green). **d**. Quantification of cluster formation in scrambled or glypican-4 knockdown SCs at 72 hours post-knockdown. (mean ± SEM) n=3. *p<0.05, **p<0.01, ***p<0.001. Scale bar=50µm. **e.** Representative confocal images showing scrambled or Slit2/3 knockdown cells stained with Actin (red) or immunolabelled with N-cad (green). Note stronger N-cad staining at points of cell contact. Scale bar=50µm. **f.** Quantification of **(e)** to show the percentage of SCs in clusters in scrambled or Slit2/3 knockdown cells (mean ± SEM) n=3. ***p<0.001. **g.** Quantification of CIL in scrambled treated or Slit2/3 siRNA knockdown cells (mean ± SEM) n=3. *p<0.05, **p<0.01.

Glypicans are GPI-linked proteins that have been reported to act as co-receptors in several signalling pathways including Robo, Wnt, FGF, Hedgehog, and the bone morphogenetic pathways and can modulate the accessibility of ligands but are not thought to act as ligands themselves, suggesting another signal may be required^41, 42^. Of particular interest was the Slit-Robo signalling pathway which can play a role in axonal guidance^42–47^. Previous studies have shown that of the three Slit genes, only Slit2 and Slit3 are expressed by SCs, while Robo1 has been reported to be expressed by SCs and axons^48, 49^. We confirmed the expression of Slit2 and Slit3 in SCs (Supplementary Fig. 4e), and performed knockdowns of both (Supplementary Fig. 4d). Although, we saw a small loss of CIL with each siRNA, it was not as strong as seen with Glypican-4, possibly because of compensation between the two molecules. We therefore performed a double knockdown of Slit2 and Slit3 using siRNA. Importantly, Slit2/Slit3 knockdown cells behaved similarly to Glypican-4 knockdown cells and did not repulse upon contact, but formed cell clusters with an increase in N-cadherin-mediated junctions compared to the control (Fig. 4e-f). Moreover, similarly to the Glypican-4 knockdown cells, the cells failed to show CIL within the clusters (Fig. 4g). Together these results show that Slit2 and Slit3 mediate CIL between SCs.

### N-cadherin is required for the trafficking of Slit2/Slit3 to the cell-surface

We have shown that both N-cadherin and Slit2/Slit3 are required for CIL between SCs. We hypothesised that this could be due to the loss of the repulsive Slit2/Slit3 signal in cells lacking N-cadherin. However, following N-cadherin knockdown, little change was detected in total Slit2 and Slit3 protein levels (Supplementary Fig. 5a) and RNA levels were not affected (Supplementary Fig. 5b) indicating a post-translational mechanism was involved. Co-immunoprecipitation experiments showed that Slit2 could be pulled down by N-cadherin (Supplementary Fig. 5c) and live-imaging of SCs expressing tagged N-cadherin showed that N-cadherin is a highly dynamic protein arriving in waves towards the cell’s moving front suggesting that perhaps a pool of N-cadherin was involved in the presentation of Slit2 and Slit3 at the cell-surface (Supplementary Fig. 5d). To test this we performed immunostaining of control and N-cadherin knockdown cells. Confocal images of scrambled-treated SCs showed that Slit2 and Slit3 were localised at the membrane, and could be detected on the surface of the lamellapodia (Fig. 5a-d). Moreover, both ligands could be detected in numerous vesicles throughout the cytoplasm. In contrast, in N-cadherin knockdown cells Slit2 and Slit3 were no longer observable in cytoplasmic vesicles or at the cell-surface but rather were sequestered in the perinuclear region (Fig. 5a, c, quantified in Fig. 5b, d). Importantly, the trafficking of Slit2 and Slit3 could be restored by expression of the extracellular domain of N-cadherin (Fig. 5e, g, quantified in Fig. 5f, h). Together these results show that a pool of N-cadherin, via the extracellular domain, is required to traffic Slit2 and Slit 3 to the cell-surface where they initiate CIL.

**Figure 5.**
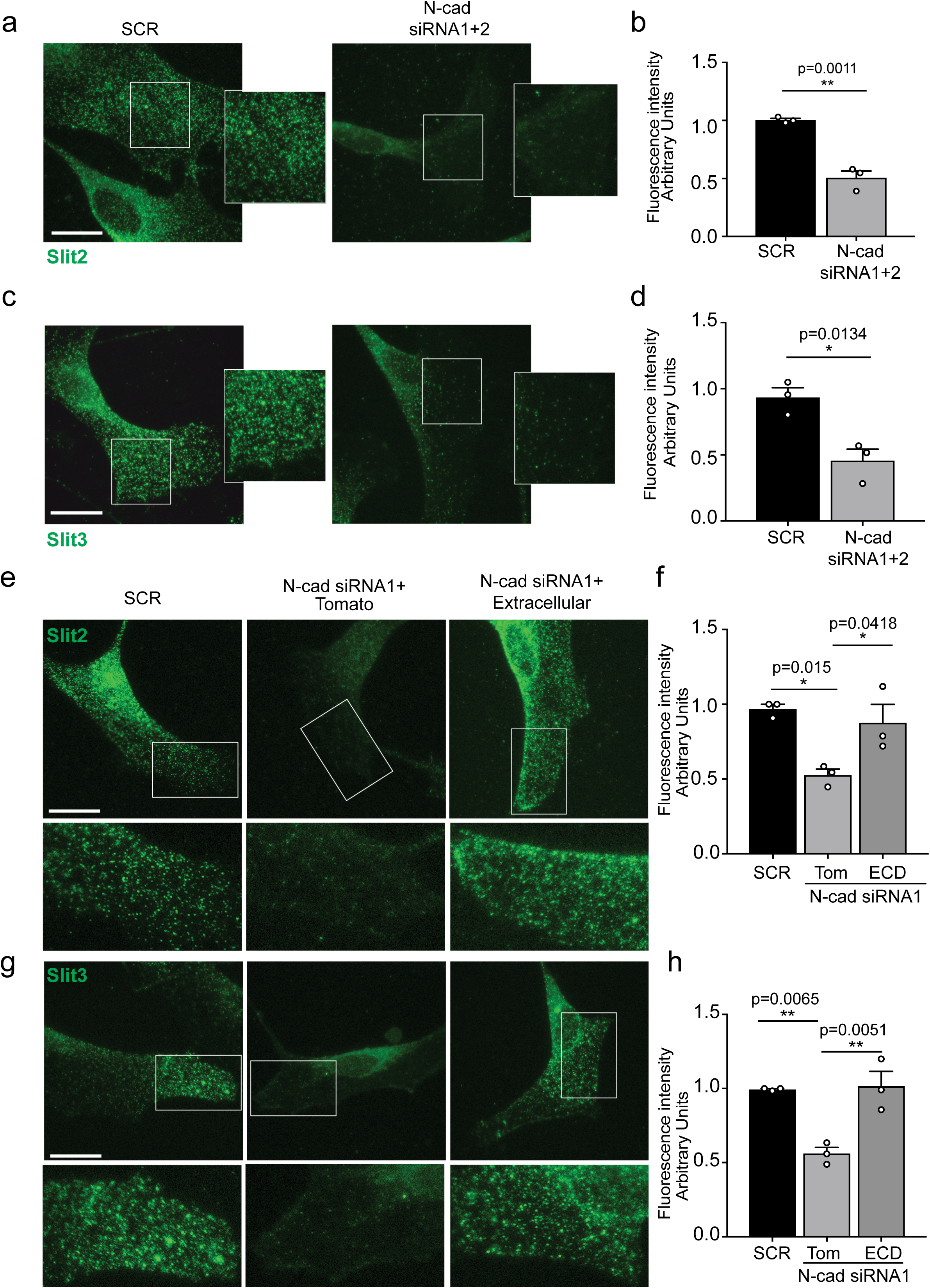
N-cadherin is required for the trafficking of Slit2/Slit3 to the cell-surface. **a, c.** Representative confocal images of scrambled or N-cad knockdown SCs. Cells were labelled with antibodies to **(a-b)** Slit2 or **(c-d)** Slit3 (green). **b, d.** Quantification of Slit2 and Slit3 levels, respectively, in cell protrusions as indicated by the boxes in **(a, c)** (mean ±SEM) n=3. *p<0.05, **p<0.01. Scale bar=15µm. **e.** Representative confocal images of N-cad knockdown SCs (siRNA1) expression ECD-tagged with tomato, or tomato control stained with Slit2 (green). **f.** Quantification of Slit2 levels in **(e)** (mean ±SEM) n=3. *p<0.05. Scale bar=15µm. **g.** Representative confocal images of N-cad knockdown SCs (siRNA1) expression ECD-tagged with tomato, or tomato control stained with Slit3 (green). **h.** Quantification of Slit3 levels in **(g)** (mean ±SEM) n=3. **p<0.01. Scale bar=15µm.

**Figure 6.**
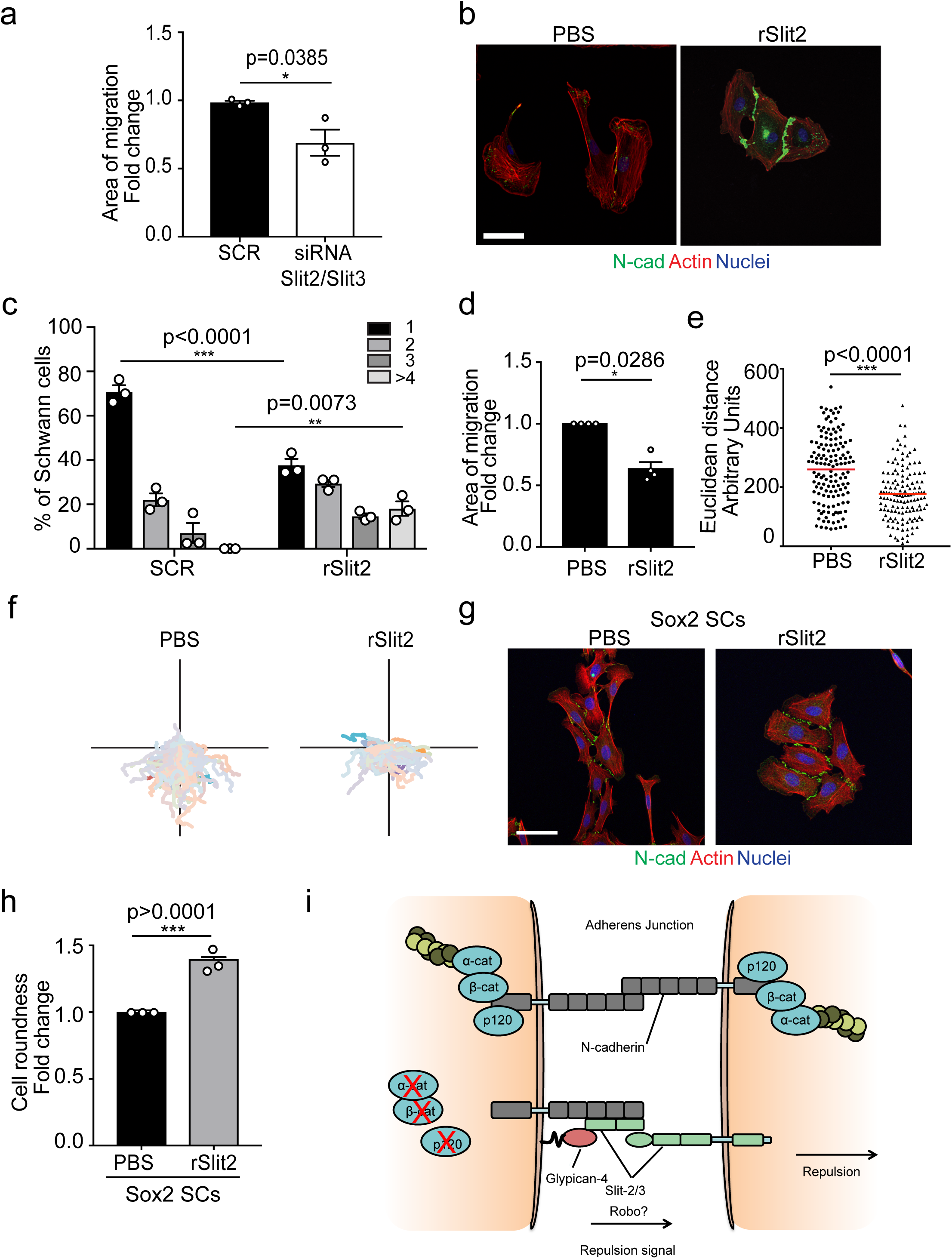
Slit is required for the efficient collective migration of Schwann cells. **a.** Quantification of the area of migration in scrambled and Slit2/3 knockdown SCs. Area of migration is expressed as fold change relative to scrambled cells (mean ± SEM) n=3. *p<0.05. **b**. Representative confocal images of SCs treated with recombinant-Slit2 (rSlit2) or PBS and stained for Actin (red), N-cad (green) and Nuclei (blue). n=3. Scale=50µM. **c**. Quantification of SC clusters from **(b)** (mean ±SEM). **p<0.01, n=3. ***p<0.001. **d.** Graph showing the area of migration covered of the wound-healing assay (mean ± SEM) n=3. *p<0.05. **e**. Graph showing the Euclidean distance (shortest distance travelled) for cells in **(d)** (PBS n=141, rSlit2 n=138). The red line denotes the mean. ***p<0.001. **f.** Graph showing tracks of wound-healing assay in **(d)** n=3. **g.** Representative confocal images of SOX2 SC clusters treated with Shield for 24 hours, +/- PBS or rSlit2 and stained with N-cad (green), Actin (red) and Nuclei (blue). Scale bar=50µM. **h.** Quantification of cell shape of SOX2 SC clusters treated with PBS or rSlit2 (mean ±SEM) n=3. ***p<0.001 **i.** Schematic representation of our proposed model for CIL in SCs.

### Slit is required for the efficient collective migration of Schwann cells

CIL can provide an outward force in collectively migrating cells^21–23^. To test if the Slit CIL signal is important for the collective migration of SCs, we performed a wound-healing assay with SCs knocked down for Slit2 and Slit3 and found that this was sufficient to decrease the collective migration of these cells (Fig. 6a, Supplementary Fig. 6a). The ability to inhibit the CIL signal whilst maintaining cell-cell contacts has therapeutic potential in that in contrast to producing invasive cells (N-cadherin inhibition) it should result in the formation of more tightly clustered, less migratory cells. Recombinant Slit2 (rSlit2), has been reported to locally induce a repulsive signal and mediate a collapse of protrusions^48^, and thus could potentially act in a dominant-negative manner possibly by providing a uniform signal around cells. To test this, we initially analysed the effect of rSlit2 on CIL in low density cultures and found that rSlit2 appeared to act as a dominant-negative to inhibit Slit-mediated repulsion signalling, as rSlit2-treated SCs no longer repulsed each other and instead formed clusters (Fig. 6b, quantified in 6c) similar to that observed when Slit signalling was inhibited using siRNA (Fig. 4g). Next we performed a wound-healing assay in cells treated with rSlit2 and found that cells treated with rSlit2 migrated less efficiently than controls, with a slower closure of the wound (Fig. 6d). By tracking the individual cells, this behaviour was distinct from N-cadherin knockdown cells in that the cells migrated a shorter distance (Fig. 6d-f) and appeared to adhere to each other. This prevented them from migrating forward which is consistent with the loss of repulsion signal observed when Slit2 signalling is inhibited at low density (Fig. 6b-c and 4f-h). SCs migrate collectively as cords during nerve regeneration in response to a Sox2 signal that, via the formation of more stable N-cadherin junctions, is dominant over the CIL signal. Loss of the CIL signal should therefore maintain these junctions while blocking the outward force required for collective migration. Consistent with this, we found that the addition of rSlit2 to Sox2-induced SC clusters resulted in clusters with a distinct morphology that lacked polarisation away from the cluster (Fig. 6g, quantified in 6h, Supplementary Fig. 6c) suggesting a lack of repulsion between the cells^50^. Together, these data show that Slit signalling is important for the collective migration of SCs and highlights the potential of targeting CIL signals to inhibit collective cell migration (Fig. 6i).

## Discussion

SCs migrate collectively as cellular cords following the severing of a peripheral nerve, guiding re-growing axons back to their targets, and thereby promoting peripheral nerve regeneration^12-14, 51^. Collective migration is a complex and continuously adapting process that requires the incorporation of multiple and opposing intercellular signals between individual cells within the group, as well as signals from the local environment such as chemotactic factors, which may either guide and induce migration, or prevent the migrating cells from invading tissues^24, 52^. During migration intercellular-repulsive signals direct the migration of the cluster forward, whilst adhesive interactions are needed to maintain a cohesive group. How these seemingly opposing forces are co-ordinated to achieve collective migration remains mostly unclear^24, 52^.

We previously showed that, similarly to other cell-types such as astrocytes and neural crest cells^21, 31, 53^, the adherens junction molecule N-cadherin is required for collective SC migration by mediating the clustering of the SCs^12^. In this study, we show that N-cadherin is also required for the repulsion signal, which provides an outward force for collective SC migration. N-cadherin has previously been shown to mediate the repulsion signal important for the collective migration of neural crest cells^21, 22, 54^. However, despite neural crest cells and SCs belonging to the same lineage, we find that N-cadherin acts by a different mechanism to regulate CIL in SCs; CIL between neural crest cells was reported to require the trans-homodimerisation of N-cadherin that acted to inhibit local protrusion formation at cell-cell contacts via the adhesion complex^21, 22, 54^. In contrast in SCs, while cell-cell adhesion is mediated by N-cadherin homodimers linked to the actin cytoskeleton via the adherens junction complex, N-cadherin mediates CIL by a distinct mechanism. This does not require trans-homodimerisation, is independent of the adherens junction complex and only requires the extracellular-domain. Instead N-cadherin is required for the trafficking of the Slit2/Slit3 repulsion signal to the cell surface. The reasons for these differences remain unclear but could reflect that whereas SCs migrate in cord-like structures, neural crest cells migrate in much looser clusters, which may require different mechanisms to maintain an adhesive migratory cluster.

Slits are secreted axonal guidance molecules that are crucial for the development of the brain^42-44, 46, 47, 55^. However despite being secreted, accumulating evidence suggests that the secreted forms can remain associated with the plasma membrane which would allow the local cell-cell signalling required for CIL^56^; indeed, a recent report has suggested that CIL between fibroblasts involves Slit2/Robo4 signalling^57^. Here, we show that Slit2/Slit3 signalling is responsible for the homotypic CIL signal between SCs and that this signal is required for the efficient directional collective SC migration that is required for nerve repair. This is consistent with reports that Slit2, Slit3 and Robo1 are expressed by myelinating and non-myelinating SCs^48, 49^ and that a Slit2 signal is sufficient to repulse a SC in culture^48^. The ability of a secreted Slit signal to act as a repellent over both longer and short distances likely requires cell-specific mechanisms that regulate whether the Slit molecules remain tethered to the cell surface^58, 59^. Our work shows that Slit-dependent CIL between SCs requires both the extracellular domain of N-cadherin and Glypican-4 and it therefore appears likely that this complex is important for the trafficking and retention of Slit2/Slit3 at the cell-surface. This is consistent with previous reports in which Glypican and other heparan sulfate proteoglycans appear to be important for local Slit signalling^60–62^.

Further complexities of the role of CIL in cell behaviour are indicated by the distinct mechanisms used to regulate heterotypic and homotypic CIL and how these signals can change the migratory behaviour of cells. SCs show complex migratory behaviours influenced by the microenvironment; SCs alone exhibit CIL, which we now show is a N-cadherin/Glypican4/Slit2/Slit3 dependent process that repulse SCs from each other, which is perhaps important for the distribution of SCs along axons. Following an injury however, SCs come into contact with fibroblasts at the injury site and the fibroblasts repulse SCs by an ephrinB/EphB2–dependent signal^12^. Importantly however, this signal also changes the behaviour of N-cadherin resulting in the formation of more stable N-cadherin junctions that act dominantly over the CIL signal resulting in the formation of cell cords. However, the Slit2/Slit3-dependent repulsion signal remains active within the cords, providing an outward force to drive the collective migration of these cells. While both N-cadherin and Slit2/Slit3 are important for the collective migration of cells, inhibition of these proteins has very distinct effects on the migration of SCs. So whereas loss of N-cadherin results in the production of invasive, highly migratory cells that lack collective behaviour, inhibition of Slit2/Slit3 results in the formation of tighter clusters of cells associated with stronger N-cadherin junctions in which migration is inhibited (Supplementary Fig.S6d). This might suggest targeting of CIL signals would be a promising approach for inhibiting collective migration. Consistent with this, we showed that exogenous recombinant Slit2 acts to inhibit SC CIL and inhibits the collective migration of these cells. Collective migration has been shown to be crucial for tumour cell migration with a role proposed for CIL in this process^50, 63–66^, identification of the CIL signals of distinct tumour types may therefore be a promising approach to identify new targets for inhibiting the invasion and spread of tumours.

## Acknowledgements

This work was supported by a programme grant from Cancer Research UK (C378/A4308) and core support by MRC funding to the MRC LMCB University Unit at UCL, award code MC_U12266B. We would like to thank Lucie Van Emmenis, Liza Malong and the rest of the Lloyd lab for useful discussions.

## Author information

The authors declare no competing financial interests. Correspondence and requests for materials should be addressed to A.C.L.

## Materials and Methods

### Cell culture

Primary rat Schwann cells (SCs) were extracted from sciatic nerves of postnatal 7 days Sprague-Dawley rats as previously described ^67^. SCs were cultured on PLL coated dishes in low culture DMEM (Lonza) supplemented with 3% FBS (BioSera), 1 µM Forskolin (Abcam), 200mM L-Glutamine (Gibco), GGF, 100µg/mL kanamycin (Gibco) and 800µg/mL gentamicin (Gibco), and maintained in 10% CO_2_ and 20% humidity at 37ºC. HEK293T cells were cultured in DMEM (lonza) supplemented with 10% FBS and 200mM L-Glutamine (Gibco).

Sox2 overexpressing SCs were produced using the Retro X ProteoTuner Shield system (Clontech). The Retro X ProteoTuner retroviral vector encodes a 12kDa FKBP destabilization domain (DD). The DD is expressed as an N-terminal tag on the protein of interest and causes rapid degradation of the protein to which it is fused. Accumulation of the DD tagged protein of interest can be rapidly induced by addition of Shield1 stabilizing ligand to the media, which prevents proteasomal degradation of the protein.

For analysis of Sox2 clusters, Shield1 (200nM, Takara Bio) was added to Sox2 overexpressing SC or ProteoTuner SC controls to induce clustering and cells were fixed 24h following treatment.

### Constructs

N-cadherin full length, the extracellular and intracellular domain of N-cadherin tagged with ptdTomato on the C-terminus, were a kind gift from Prof. S. Yamada ^37^. The myc-tagged slit-2 construct was gifted by Dr. V. Castellani ^68^.

### Antibodies

Primary antibodies were used against: N-cadherin (BD transduction); Alphα-catenin (Sigma C2081); β-catenin (BD transductions 610920); p120-catenin (BD Transduction 61034); ERK1/2 (Sigma M5670), mCherry (Abcam ab183628; Westernblotting); mCherry (Life technologies M11217; Immunoprecipitation), AKT 1/2/3 (Santa Cruz), Slit2 (Abcam ab134166; Westernblotting), Slit2 (Thermo Fisher Scientific PA531133; Immunofluorescence), Slit3 (Sigma SAB2104337; Immunofluorescence) Slit3 (R&D Systems AF3629; Westernblotting), Myc (Merck Millipore 05-724). Alexa-Fluor secondary antibodies were obtained from Invitrogen. Horseradish peroxidase (HRP)-linked antibodies were obtained from GE-healthcare.

### Short interference RNA

All short interference RNA (siRNA) were supplied by Qiagen, scrambled siRNA (Qiagen) or all star scrambled siRNA (Qiagen) was used a control. siRNA knockdown was performed according to the manufacturer’s protocol and optimised for SCs. In brief, 10^5^ cells Schwann cells were on 6 well plates. The following day siRNA or scrambled siRNA were mixed in plain DMEM with Hiperfect (Qiagen) and incubated for 10 min at room temperature (RT) to allow complexes to form. Complexes were added to SCs for 16-18 hours, washed once with normal SC medium and harvested or seeded for further experiments as appropriate.

αE-catenin siRNA1 AAGAACGCCTGGAAAGCATAA

αE-catenin siRNA2 CAACCGGGACTTGATATACAA

Cadherin-2 siRNA1 TCCCAACATGTTTACAATCAA

Cadherin-2 siRNA2 CAGTATACGTTAATAATTCAA

p120-catenin siRNA1 AGGTCAGATCGTGGAAACCTA

p120-catenin siRNA2 ATGCTCGGAACAACAAAGAGTTAA

Glypican-4 siRNA1 CCGACTGGTTACTGATGTCAA

Glypican-4 siRNA2 CGGTGTAGTTACAGAACTGTA

Slit2 siRNA1 ATCAATATTGATGATTGCGAA

Slit2 siRNA2 GACGACTAGACCGTAGTAATA

Slit3 siRNA1 AACGGCGGTGCCCAAAGAATT

Slit3 siRNA2 ATCGTGGAAATACGCCTAGAA

### Mutagenesis

To disrupt the annealing of N-cad siRNA to the tomato-tagged constructs N-cad full-length, or the extracellular domain of N-cad, the NEB Q5 Site-Directed Mutagenesis Kit (NEB) was used according to manufacturers protocol to introduce silent mutations in the N-cad siRNA1 targeting sequence, by mutating all four codons of the target sequence.

The following primers were used:

Forward primer 5’-CACGATAAACAATGAGACTGGGGACATC-3’

Reverse primer 5’-AACATATTGGGTGAAGGTGTGCTGGG-3’

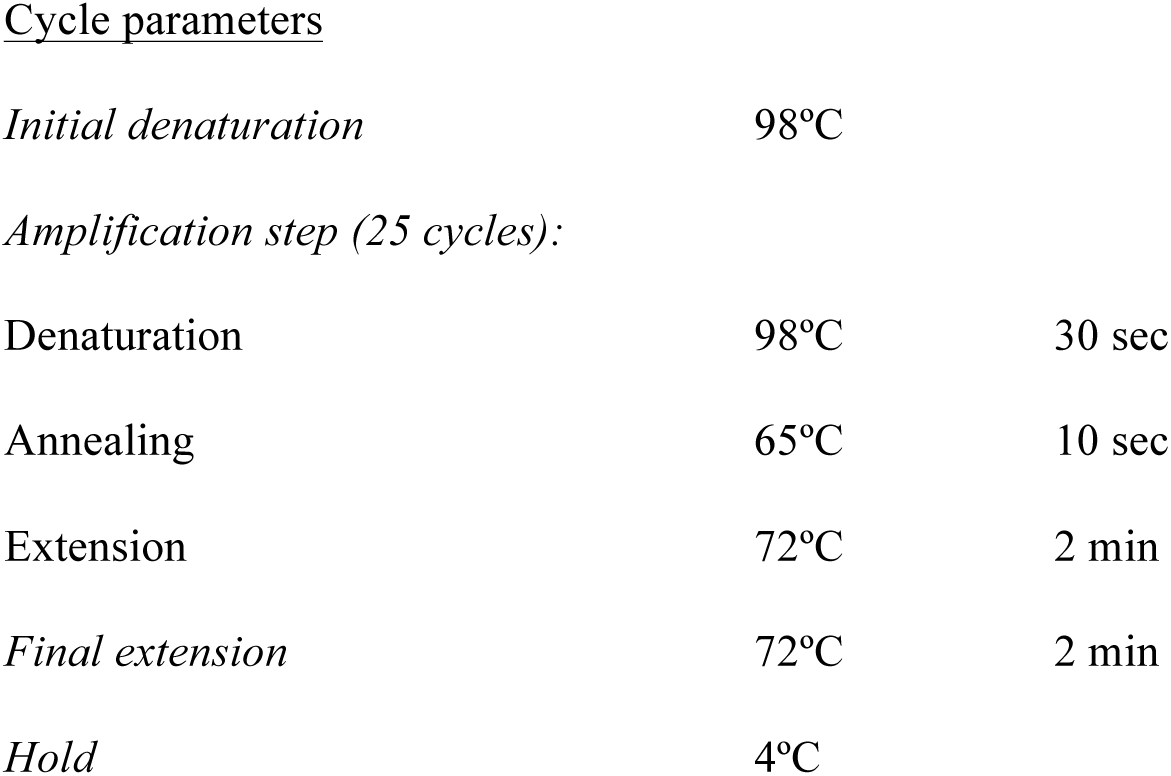

### Transfection of HEK293T cells

DNA was transfected using Attractene protocol (Qiagen). Briefly, HEK293T cells were seeded onto 60mm plates 24 hours prior to transfection. 270ng Myc-tagged Slit2 and 270 ng of Tomato-tagged constructs of interest and 1.25µg carrier vector DNA was incubated with Attractene in DMEM for 15 min at RT to allow complexes to form. Complexes were incubated for 2 hours at 37ºC.

### Contact inhibition of Locomotion assays

For repulsion assays 8×10^3^ or 4×10^3^ Scrambled treated or siRNA-treated SCs were seeded onto PLL and laminin coated 6 well plates or 12 well plates respectively. Cells were allowed to adhere for 6 to 18 hours hours before time-lapse microscopy.

To analyse interactions between scrambled and N-cad knockdown cells, cells were treated with 10µM Cell Tracker Red CMTPX Dye or Green CMFDA dye (Invitrogen) for 30 min at 37°C and washed once with 3% medium. Subsequently, 1 hour after treatment 4 ×10^3^ scrambled treated cells were mixed with 4 ×10^3^ N-cad knockdown cells, incubated for at least 16 hours before starting live-imaging using time-lapse microscopy.

### Contact inhibition of Locomotion quantifications

Volocity or Fiji software was used to quantify repulsion. Single cells that were not dividing were tracked until contact with another cell and the initial response upon contact determined. Three types of events were defined: <u>Repulsion</u>, cells retract protrusions and change direction of migration; <u>No-repulsion</u>, cells interact for longer than five hours and don’t change direction of migration; <u>Invasion</u>, cell migrate on top of another cell with their protrusion and or cell bodies. To ensure a specific readout only single cell-cell interactions were quantified. Analysis was done by eye and where possible blind.

For mixing experiments, the response of scrambled-treated cells, were quantified upon contact with N-cad knockdown cells and vice versa. Similarly, for the rescue experiments only the response of N-cad knockdown cells were quantified upon contact with a cell expressing, either GFP, the full length of N-cad, the extracellular domain of N-cad or the intracellular domain of N-cad.

### Rescue

Rescue experiments were performed in a two faceted protocol, first siRNA transfection was performed using 1nM of N-cad siRNA1 as described above, incubated overnight. The following morning complexes removed and replaced with 2mL medium, and 4-5 hours later DNA was transfected using Attractene (Qiagen). Briefly, 575ng carrier vector, 25ng pBird were and 80ng plasmid DNA carrying either N-cad full length, the extracellular domain or the intracellular domain of N-cad were mixed in plain DMEM and incubated with Attractene for 15 min at RT to allow complexes to form. Complexes were dropped onto scrambled-treated or N-cad knockdown cells and were incubated for 2 hours at 37ºC. pBird was used as control for transfection efficiency. The following day cells were seeded for repulsion assays and for immunofluorescence. Time-lapse was started approximately 6 hours after seeding and were imaged for 24 hours.

For immunofluorescence of Slit-2 and Slit-3, 600ng carrier vector and 80ng Extracellular domain of N-cad-tagged with tomato, or tomato control was used.

### Recombinant Slit2 treatment

Cells for cell clustering assay or chamber assay, were pre-treated with 2ug/mL recombinant mouse Slit2 protein (gln26-gln900) (rSlit2) (R&D system, 5444-SL-050) or PBS for at least 18 hours before removing inserts, and further incubated with 2ug/mL rSlit-2 or PBS for 24 hours while imaged using time-lapse microscopy.

### Scratch assays, invasion assays and wound healing assays

For the scratch assays cells were seeded on laminin-coated plates for a siRNA knockdown at 1×10^5^, cells. siRNA transfection was performed as described above and the cells were grown to confluence. At 48 hours after treatment a scratch was induced with a sterile tip. The cells were gently washed twice with medium to remove any debris and imaged using time-lapse microscopy for 24 hours. Cells at the leading edge were imaged.

For invasion assays, 1.5×10^4^ scrambled-treated cells or N-cadherin knockdown cells (24hours after knockdown) were seeded in the compartments of the 2 well cell-culture inserts (Ibidi), and allowed to adhere for 18 hours, before treating the cells in the two different compartments with either 10µM CellTracker-Green or Cell Tracker Red CMTPX Dye, for 30 min. Subsequently, the chamber was removed and invasion was observed.

For wound healing assays using chambers, 1.5×10^4^ cells were seeded in both compartments of two well cell-culture inserts (Ibidi), and allowed to adhere for 24 hours. Cells were treated with rSlit-2 as described above.

### Cell tracking

Cells were tracked by their nucleus using volocity software or the manual cell tracking plugin in FIJI for 8 - 12 hours, except for dividing cells. Velocity and the directionality were then measured from these tracks using macro plugin in excel as described in ^69^.

### Time Lapse Microscopy

Live-imaging was performed at 10x or 20x using a Zeiss Axiovert 200M microscope equipped with a Hamamatsu Orca AG camera controlled by Volocity software (Improvision) and an environmental chamber which maintained the temperature at 37ºC and provided a humidified stream of 5% or 10% CO_2_ in air. Images were taken every 10 minutes for 24 or 48 hours 72 hours. Live imaging of tomato tagged N-cadherin was performed on the Ultraview Vox spinning disc confocal microcope, and maintained at 37ºC with 10% CO_2_. Images were of two cells interacting were acquired every 20 seconds and analysed using Volocity software (Improvision).

### Cell clustering assays

Cell clustering assay was perfomed as described in ^*12*^. ^*12*^In brief, 3.5 x 10^3^ cells treated with Glypican or Slit2/3 siRNA were seeded onto coverslips 72 hours and 96 hours after knockdown fixed and stained and clusters were quantified 24 hours after seeding the cells. For rSlit-2 treated cells, 3.5 x 10^3^ cells were seeded onto 12-well plate and rSlit-2 treatment was performed as described above, clusters were quantified after 36 hours of treatment. The following clusters were defined, single cells, 2, 3, or clusters existing of >4 cells.

### Immunofluorescence

Cells grown on glass coverslips were fixed in 4% paraformaldehyde (PFA) supplemented with 1mM CaCl_2_ and 0.5mM MgCl_2_ to prevent disruption of the Ca dependent complexes such as N-cadherin and the cytoskeleton for 10 minutes at RT, permeabilised with 0.3% Triton in PBS for 10 minutes at RT and blocked with 3% Bovine serum albumin (BSA) in PBS for 1 hour at RT. Coverslips were incubated with primary antibodies overnight at 4ºC. The following day, coverslips were washed with PBS, incubated with secondary antibodies conjugated to Alexa Fluor for one hour at RT and washed with PBS before mounting onto microscope slides using fluoromount-g (Southern Biotech).

For labelling of Slit2 and Slit3, cells were incubated for 10min with CellMask deep red (lifetechnologies) for 10 min and fixed in 2% PFA supplemented with 1mM CaCl_2_ and 0.5mM MgCl_2_ and blocked 3% Bovine serum albumin (BSA). All remaining steps were performed as described above.

All images were acquired using an inverted Leica TCS SPE confocal microscope. All image processing and analysis was performed using FIJI software. For each image, a projection of the z-stacks was made using Fiji software using the maximal intensity.

### Quantification of immunofluorescence

To quantify Slit3 and Slit2 immunolabelling, the outline of the cells was drawn, using the free hand drawing tool in Fiji and measuring the fluorescence intensity. The nucleus and perinuclear area, indicated by the brighter staining, was excluded from the quantification. The corrected total cell fluorescence was calculated using this formula: integrated-(area selected cells – mean background readings).

The shape of individual SCs in clusters was calculated using the equation 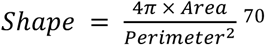 where the area and perimeter of each cell was estimated by tracing their outline using FiJi from images taken at 20x magnification. Groups of SCs were considered to be in clusters if they appeared to be flatter than normal and showed an abundance of N-cadherin staining at the cell-cell junctions.

In order to calculate the area of collective migration from chamber assays, the area of cells was drawn using FiJi at 0h and 6h or 24h on stills from time-lapse microscopy. The area migrated was calculated using this formula: Area Migrated=Area 6h-Area 0h and expressed as area fold migration change relative to PBS controls.

### Protein Analysis

Protein extraction and quantification

For protein extraction cell media was aspirated from cells which were then washed on ice with PBS, followed by snap freezing at −80°C to break membranes and harvested in 100-150µL RIPA buffer (1% Triton X-100, 0.5% sodium deoxycholate, 50mM Tris pH7.5, 100mM NaCl, 1mM EGTA pH8, 20mM NaF, 100µg/ml PMSF, 15µg/ml aprotonin, 1mM Na3VO4, 1/100 protease inhbitior cocktail). Cells were then lysed on ice for 30 minutes, vortexed every 10 minutes and homogenised using a 26 gauge needle (Beckton Dickinson). Cell debris were pelleted by spinning down at 750g for 5 min at 4ºC, the supernatant was collected and quantified using BCA assay (Pierce, Thermo Scientific).

### Western Blotting

Western blotting was performed using Hoefer Scientific Instrument apparatus and Biorad western blot electrophoresis system. 20 to 30µg of proteins were resolved using a Sodium Dodecyl Sulfate - polyacrylamide gel electrophoresis (SDS-PAGE). Protein was transferred onto nitrocellulose membrane (Millipore-Immobilon), which was subsequently blocked for 1 hour at RT using 5% milk-TBST. The membrane was incubated with primary antibodies overnight at 4ºC. The following day, the membrane was washed three times with TBST, followed by incubation with the appropriate secondary antibody conjugated to Horseradish peroxidase (HRP). Subsequently, membranes were washed three times with TBS-T before detection of proteins of interest with Pierce-ECL western blot substrate (Thermo Scientific) or Luminata Crescendo Western HRP substrate (EMD-Millipore) on the Imagequant LAS 4000.

### Co-immunoprecipitation (Co-IP)

SCs were seeded at 1.2×10^6^ onto 15cm dishes onto 4 plates and grown to sub-confluence. Cells were scraped in NP40 buffer (50mM Tris pH 7.5, 150mM NaCl, 1% NP40 supplemented with 1/100 protease cocktail inhibitor (Sigma), 1/100 phosphatase inhibitor cocktail 2 and 3 (Sigma)), lysed on ice for 30 minutes. Debris was pelleted, by centrifuging at 750g for 5 min at 4ºC and the supernatant was collected into a fresh tube. Protein concentration was then quantified using BCA assay. All subsequent steps were performed at 4ºC. Approximately 1mg protein was pre-cleared using 10µL of 50% protein G Beads (GE healthcare) by rotating for 15 minutes. Beads were collected and discarded by centrifuging at 750g for 1 minute. Pre-cleared supernatant was incubated with primary antibodies targeting N-cad (4ug), Tomato using 7ug of anti-mcherry (life technologieS) or mouse or Rat IgG as controls respectively (Santa Cruz) for 2 hours by rotating. The antibody-protein complexes were pulled down by rotating the mixture with 50µL of 50% Protein G beads for 1 hour. Beads were washed four times with 500µL NP40 buffer and were collected by centrifuging at 750g for 1 minute and transferred to a clean tube. Finally, the beads were resuspended in 30 µL Leamlli buffer and boiled for 10 minutes at 95C.

### RNA extraction and qPCR

RNA extraction was performed using Tri-Reagent according to manufactures protocol. RNA concentration was determined using Nanodrop and 500-1000ng of RNA was used to synthesize complementary DNA using Superscript II kit (Invitrogen). For RT-qPCR the MESA Blue qPCR MasterMix Plus kit was used (Eurogentec). For each gene MESA Blue was mixed with 100nM forward and reverse primers (refer to table 2.2), 23uL was pipetted in to a 96 well. 2µL of template cDNA was added to mix, ddH2O was used as a negative control.

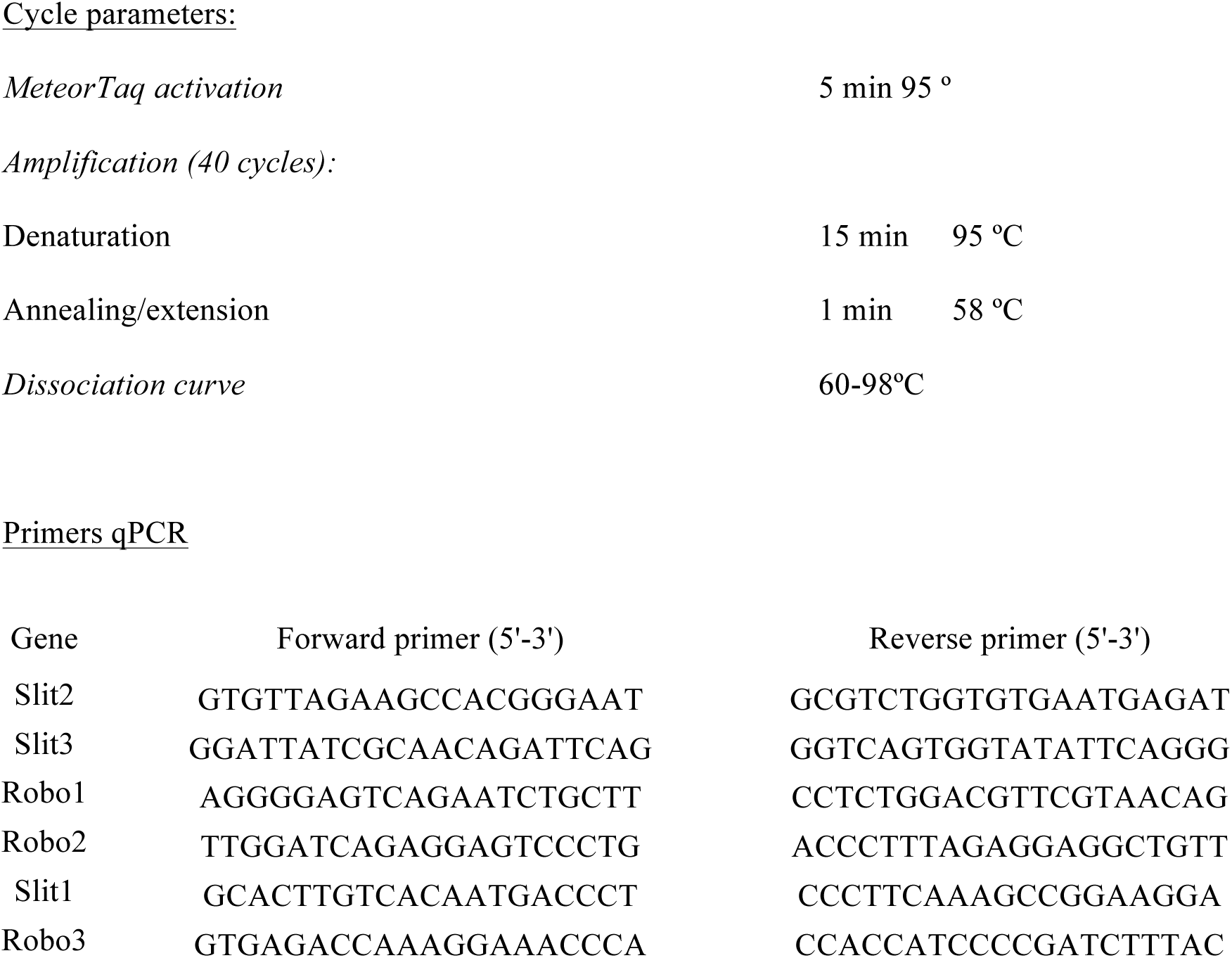

### Statistics

Statistics were performed using GraphPad (Prism) software. Data is presented as mean ±SEM, and were analysed using a one-way or two-way analysis of variance (ANOVA), followed by Tukey’s multiple comparisons tests, unpaired Student’s t-test or Mann-Whitney test, as appropriate. In all cases p *< 0.05, **<0.01, *** <0.001, n.s. = not significant.

---

**Supplementary Figure 1.**
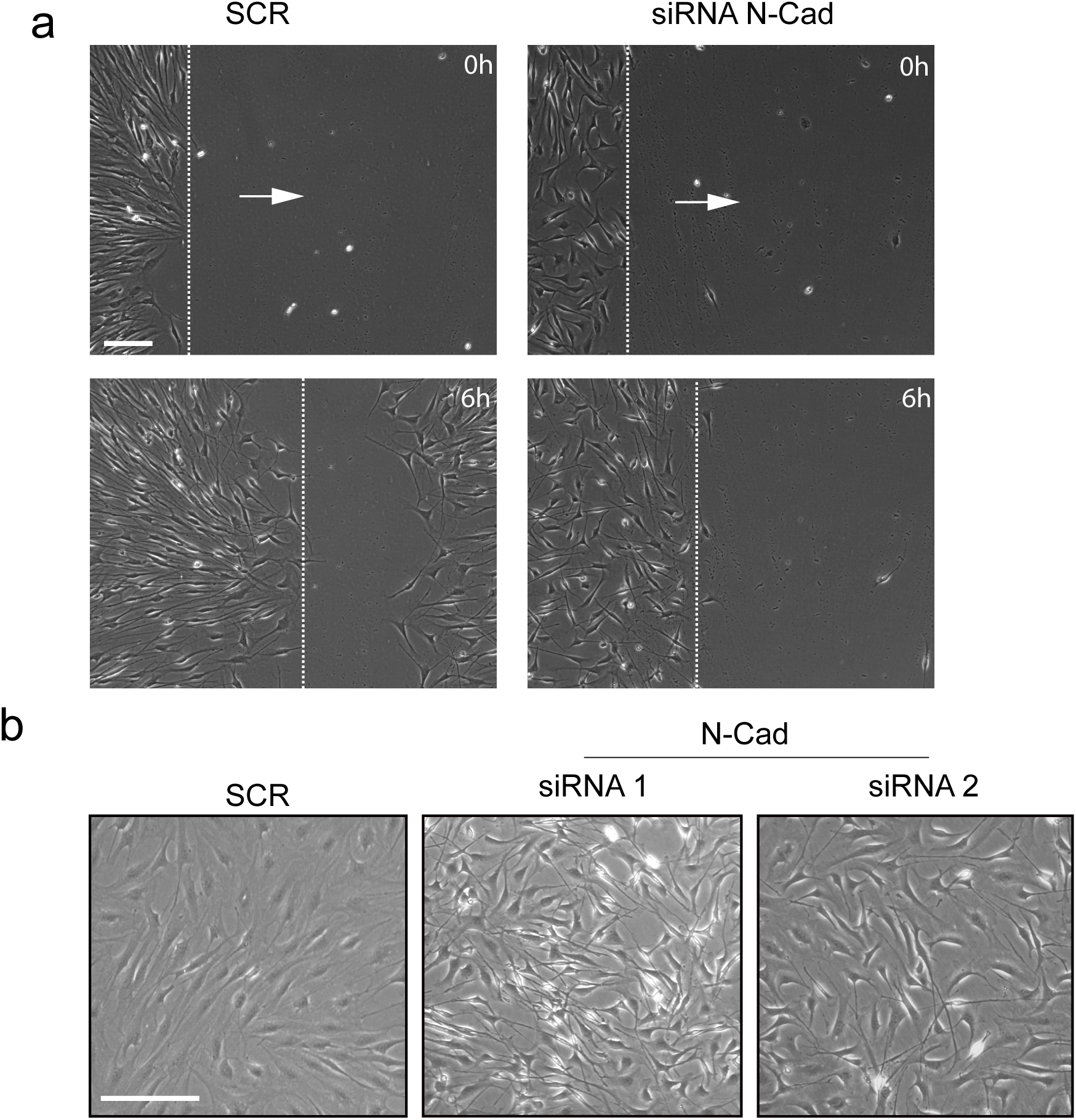
**a.** Representative still images from time-lapse microscopy of a scratch assay of scrambled treated SCs or N-cad knockdown SCs treated with siRNA1. The dashed lines indicate the leading edge of the migrating cells. Arrow indicates the direction of migration. Scale bar=100μm. **b.** Representative bright-field images of SCs treated with scrambled, or N-cad knockdown cells treated with siRNA1 or siRNA2. Note that whereas scrambled treated cells form a monolayer, N-cadherin knockdown cells grow on top of each other. Scale bar=200μm.

**Supplementary Figure 2.**
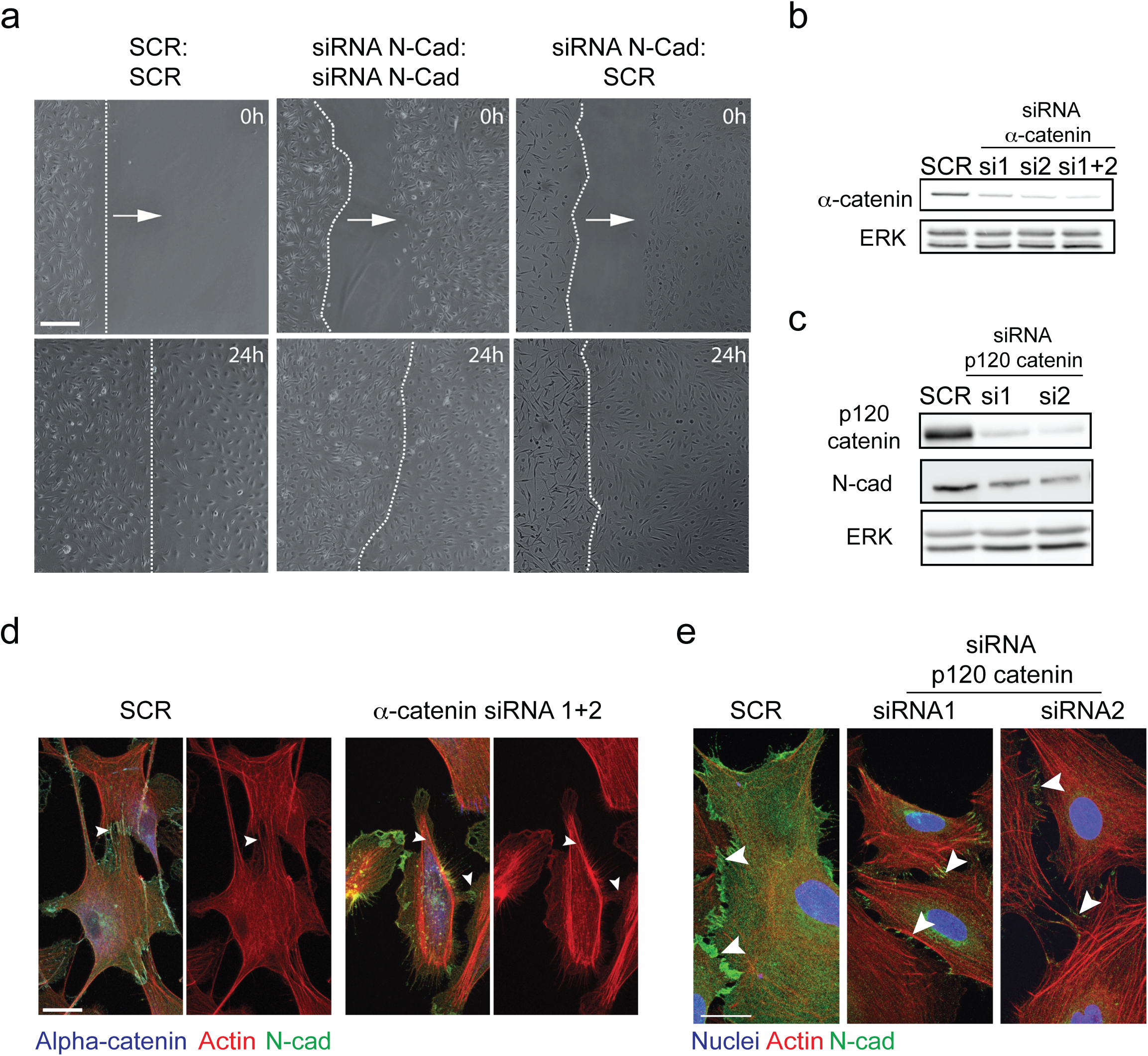
**a.** Representative image stills from time-lapse microscopy from a chamber assay of scrambled, N-cad knockd**o**wn or scrambled and N-cad knockdown SCs at 0h and 24h. The dashed lines indicate the leading edge of the migrating cells. Arrow indicates the direction of migration. Scale bar=100μm. **b.** Representative Western blot showing the efficiency of **(b)** α-catenin or **(c)** p120 catenin knockdown using two independent siRNAs compared to scrambled control at 36 hours for α-catenin and at 96 hours for p120 catenin. ERK was used a loading control. **d.** Representative confocal images of scrambled-treated and α-catenin knockdown cells at 36 hours, immunostained to detect α-catenin (blue), N-cad (green) and co-stained for f-actin (red). Note N-cad is still localised at the junctions following alpha catenin knockdown (arrows) but the actin is no longer polarised to these junctions but rather has a more cortical appearance. Scale bar=20μm. **e.** Representative confocal images of scrambled or p120-catenin knock-down cells at 96 hours, immunostained for N-cad (green), and co-stained for Actin (red) and nuclei (blue). Arrows indicate N-cad positive cell-contacts, which are fewer following p120-catenin knockdown. Scale bar=20μm.

**Supplementary Figure 3.**
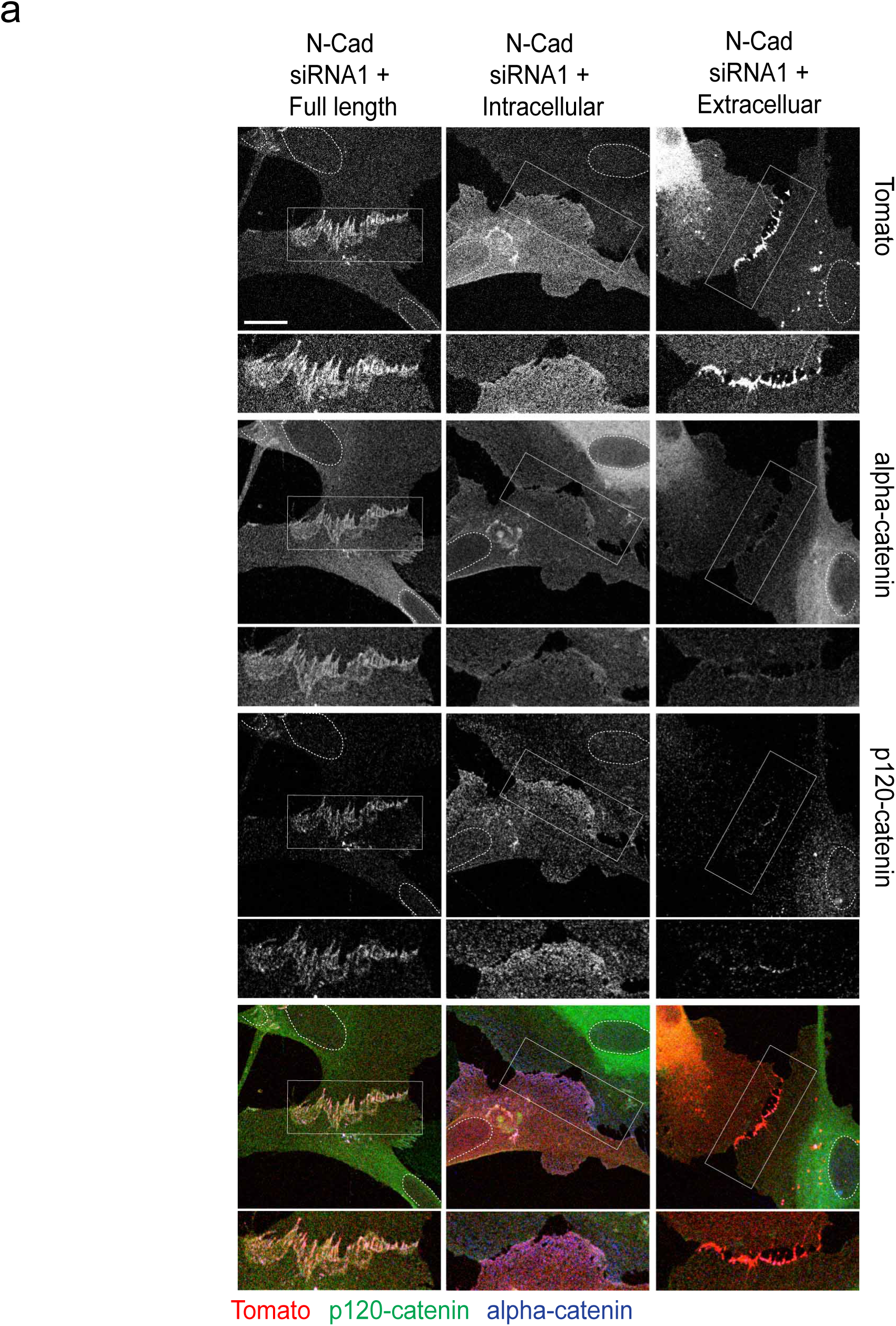
**a.** Representative confocal images of N-cad knockdown cells overexpressing ptdTomato-tagged N-cad full length (left panel), the intracellular domain (middle panel), or the extracellular domain of N-cad (right panel) (red), immunolabelled with p120-catenin (green) or α-catenin (blue). Enlarged insets show the area of cell-contacts where the constructs are localised. Scale bar=20μm.

**Supplementary Figure 4.**
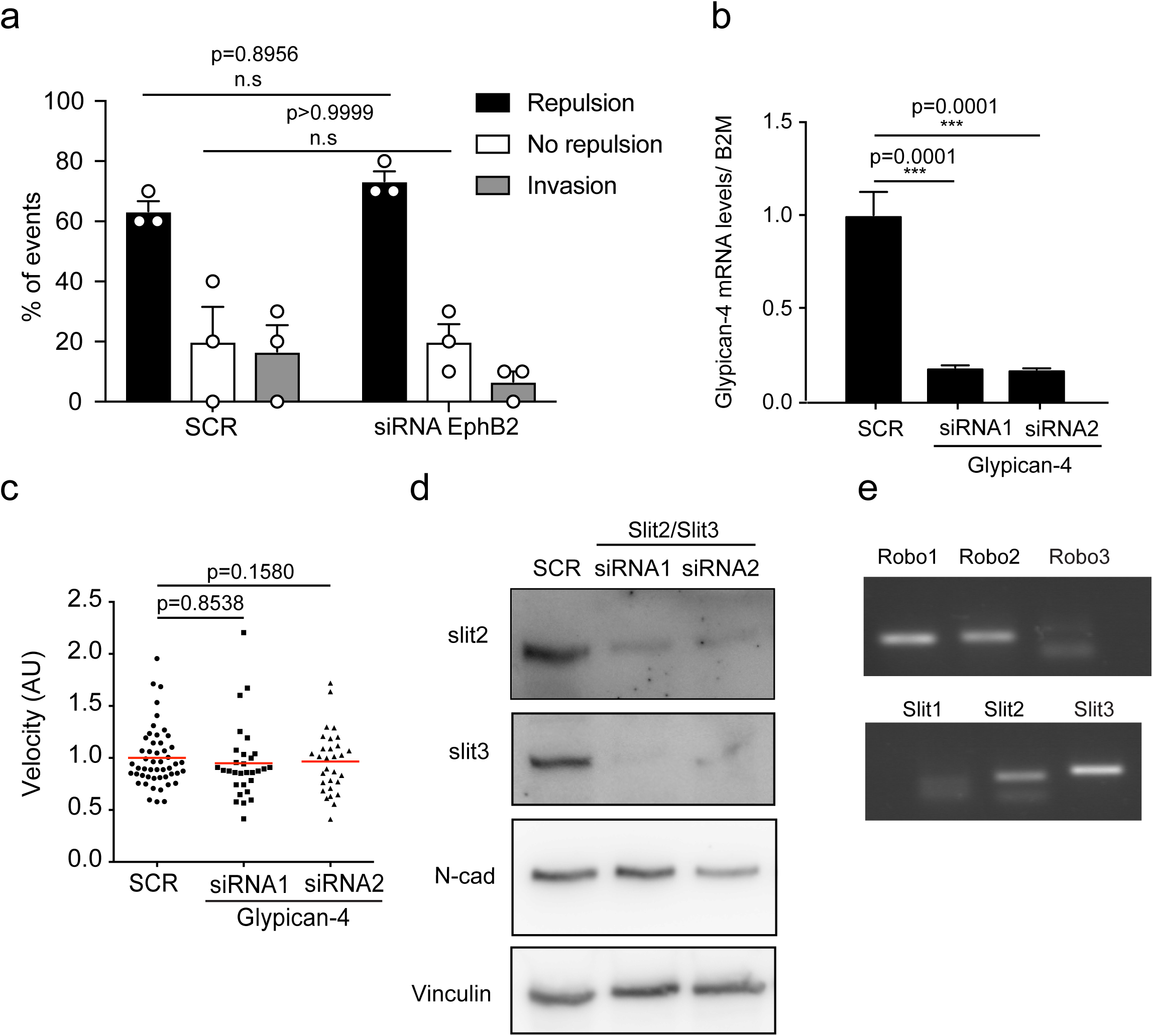
**a.** Quantification of CIL in scrambled or EphB2 knockdown SCs. (mean ±SEM). n=3 n.s=not significant **b.** Graph showing mRNA of glypican-4 mRNA levels from RT-qPCR of scrambled or glypican-4 knockdown SCs at 36 hours (mean ±SEM) n=3. ***p<0.001. **c.** Quantification of the velocity of individual cells in **Figure 4a**, red line shows the mean, n=30. **d.** Representative western blots of scrambled and slit2/3 knockdown cells, showing the knockdown efficiency and N-cad expression levels. Vinculin was used as a loading control. **e.** Gel confirming the expression of the different Slit ligand and Robo receptors in SCs.

**Supplementary Figure 5.**
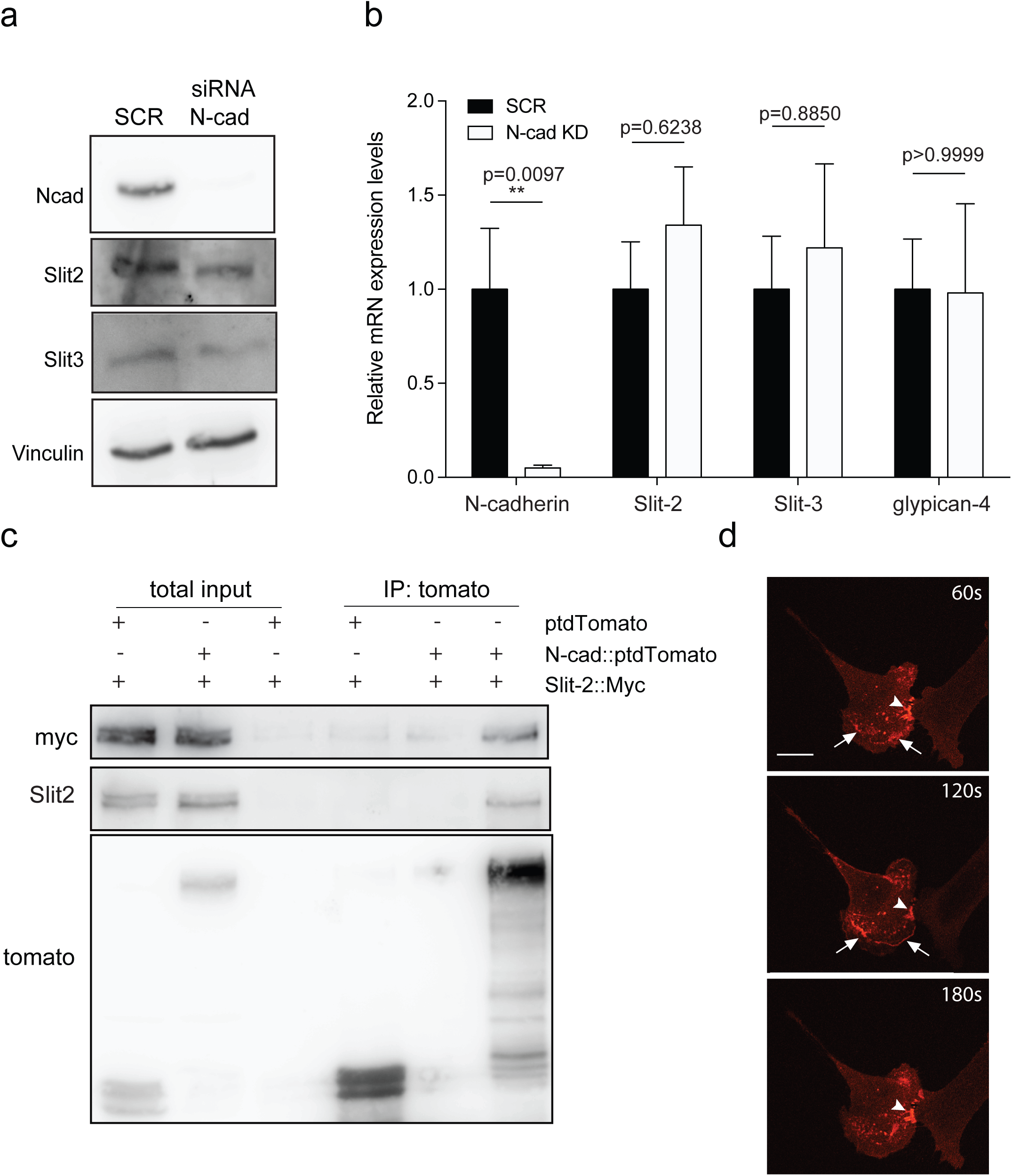
**a.** Representative western blot of scrambled and N-cad knockdown cells, probed for Slit2, Slit3 and N-cad. Vinculin was used as a loading control. n=3. **b.** Graph showing mRNA expression levels of Slit2, Slit3 and glypican-4 in N-cad knockdown cells (siRNA1 and siRNA2) compared to control (mean ± SEM) n=3. **c.** Representative western blot showing the co-immunoprecipitation of full-length N-cad tagged with Tomato, co-expressed with myc-tagged Slit2 in HEK cells. Blots were probed with anti-mcherry or anti-myc, (n=3). **d.** Representative stills from spinning disc confocal microscopy of SCs transfected with tomato labelled N-cad demonstrating the dynamic activity of this protein, which arrives in waves towards the moving front of the cell, and is visible at cell contact points. Arrowheads indicate N-cad at cell-cell contacts. Arrows indicate N-cad at the moving front of the SC. Scale bar=20 μm.

**Supplementary Figure 6.**
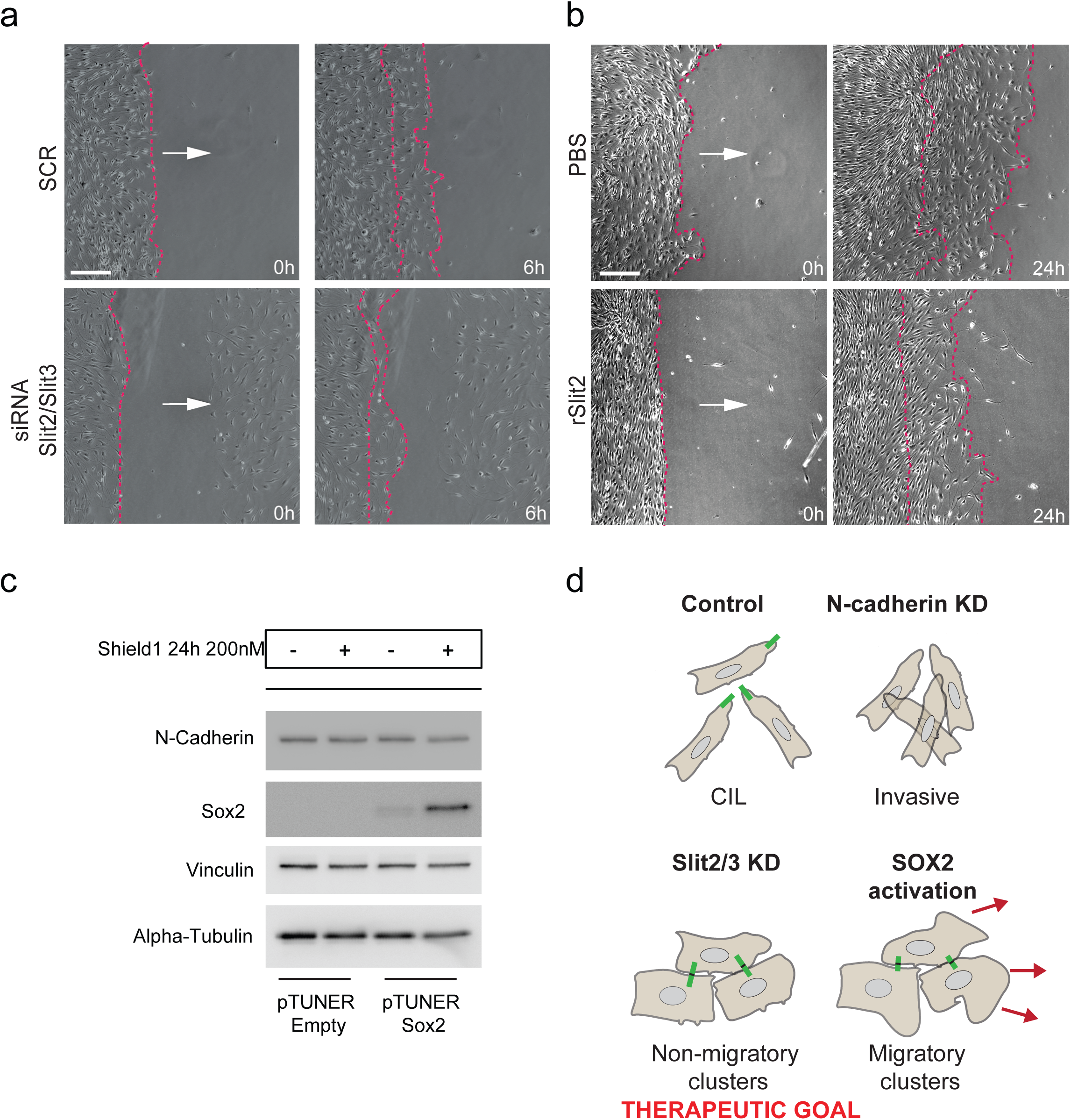
**a.**Representative stills from time-lapse microscopy of scrambled or Slit2/3 knockdown SCs seeded in chambers at 0 and 6 hours, as quantified in **Figure 6a**. (n=3) The dashed lines indicate the leading edge of the migrating cells. Arrow indicates the direction of migration. Scale bar=50 μM. **b.** Representative time-lapse microscopy images of the distance migrated (**Figure 6d**) of SCs seeded in chambers and treated with 2ug/mL recombinant-Slit2 (rSlit2) or PBS (n=3). The dashed lines indicate the leading edge of the migrating cells. Arrow indicates the direction of migration. Scale bar=50 μM. **c.** Representative western blot showing pTuner empty vector SCs or pTuner Sox2 SCs response to Shield1 treatment at 24 hours. Note that high Sox2 expression is only evident in the Shield treated pTuner Sox2 SCs at 24 hours. **d.** Schematic illustration; in control conditions SCs exhibit CIL, whereas upon knockdown of N-cadherin SCs become invasive. Knockdown or inhibition of Slit2/3 inhibits CIL resulting in non-migratory clusters. In contrast, SCs in which Sox2 is activated maintain CIL signals, which drive their collective migration. Inhibition of these CIL signals may be beneficial to prevent collective cell migration in pathological conditions such as cancer.

